# Coordination between ESCRT function and Rab conversion during endosome maturation

**DOI:** 10.1101/2024.05.14.594104

**Authors:** Daniel P. Ott, Samit Desai, Jachen A. Solinger, Andres Kaech, Anne Spang

## Abstract

The endosomal pathway is essential for regulating cell signaling and cellular homeostasis. Rab5-positive early endosomes receive proteins from the plasma membrane. Dependent on a ubiquitin mark on the protein, they will be either recycled or sorted into intraluminal vesicles (ILVs) by ESCRT. During endosome maturation Rab5 will be replaced by Rab7 on endosomes that are able to fuse with lysosomes to form endolysosomes. Whether ESCRT-driven ILV formation and Rab5-to-Rab7 conversion are coordinated remains unknown. Here we show that loss of early ESCRTs leads to enlarged Rab5-positive endosomes and prohibit Rab conversion. Reduction of ubiquitinated cargo alleviated this phenotype. Moreover, ubiquitinated proteins on the endosomal limiting membrane prevented the displacement of the Rab5GEF RABX-5 by the Rab7GEF SAND-1/CCZ-1. Overexpression of Rab7 could partially overcome this block, even in the absence of SAND-1, whereby the HOPS complex presumably acts as Rab7GEF. Our data reveal a hierarchy of events in which cargo corralling by ESCRTs is upstream of Rab conversion and thereby ESCRT-0 and ubiquitinated cargo could act as timers determining the onset of Rab conversion.

## Introduction

Cells communicate via exo- and endocytosis with their environment. These processes are also essential for the regulation of nutrient uptake and membrane domain formation at the plasma membrane (Dixson *et al*, 2023; Palm & Thompson, 2017; Rahmani *et al*, 2019; Sigismund *et al*, 2012). Once material has been taken up from the plasma membrane through the formation of endocytic vesicles, these vesicles can undergo kiss-and-run on sorting endosomes for fast recycling of proteins to the plasma membrane and eventually fuse with early endosomes (Solinger *et al*, 2020; Solinger *et al*, 2022; Solinger & Spang, 2022). The early endosome will become a sorting/maturing endosome to recycle proteins to the plasma membrane and the Golgi apparatus and to sort proteins into intraluminal vesicles (ILVs) for degradation in endolysosomes/lysosomes. Over time the endosome will, thus, mature from an early to a late endosome. During the course of maturation other processes besides recycling and sorting have to take place. Rab5 will be replaced by Rab7, and phosphatidylinositol 3-phosphate (PI3P) by phosphatidylinositol 3,5-bisphosphate (PI3,5P_2_) (Podinovskaia & Spang, 2018; Poteryaev *et al*, 2010; Wallroth & Haucke, 2018). In addition, endosomes need to acidify before fusion with lysosomes, and in many cells, endosomes move towards the cell center (Hu *et al*, 2015; Podinovskaia *et al*, 2021; Podinovskaia & Spang, 2018). This complex maturation process from early to late endosomes begs the question about the coordination between the individual processes. We have shown previously that while Rab11-dependent recycling can happen at any stage during endosome maturation, cross-talk between Rab conversion and acidification exists and Rab conversion has to be initiated before endosomes can fully acidify (Podinovskaia *et al*., 2021).

Whether Rab conversion and ILV formation are coordinated and if so, how remains unknown. Rab conversion is thought to be initiated by coincident detection of the Rab5GEF Rabex5 (RABX-5 in *Caenorhabditis elegans*) and increasing levels of PI3P on endosomes by the Rab7GEF Mon1/Ccz1 (Mon1 is called SAND-1 or CMON-1 in *C. elegans*) (Cabrera *et al*, 2014; Langemeyer *et al*, 2020; Poteryaev *et al*., 2010; Poteryaev *et al*, 2007). In addition, Rab5 can directly bind to Mon1/Ccz1 and modulate its activity (Borchers *et al*, 2023; Herrmann *et al*, 2023; Kinchen & Ravichandran, 2010; Langemeyer *et al*., 2020). These processes lead to the replacement of Rab5 by Rab7 on maturing endosomes (Borchers *et al*, 2021; Podinovskaia & Spang, 2018). Rab conversion takes only a few minutes in various experimental systems(Kinchen & Ravichandran, 2010; Podinovskaia *et al*., 2021; Poteryaev *et al*., 2010; Rink *et al*, 2005; Singh *et al*, 2014; Skjeldal *et al*, 2021; Yousefian *et al*, 2013). In parallel, ILV biogenesis promoted by the <u>e</u>ndosomal <u>s</u>orting <u>c</u>omplex required for transport (ESCRT) machinery has to occur (Appendix Fig. S1, Appendix Table S1). ESCRT-0 and -I are early acting ESCRT complexes that bind and select ubiquitinated cargo prone to degradation (Henne *et al*, 2011). The early ESCRTs corral the cargo towards the site of ILV formation (Cullen & Steinberg, 2018). There, cargo is deubiquitinated and transferred in to ILVs, which are formed by the late acting ESCRTs ESCRT-III and -IV, in conjunction with auxiliary factors (Clague & Urbé, 2006; Henne *et al*., 2011)(Appendix Fig. S1, Appendix Table S1). When sorting is completed the late endosome/multivesicular body (MVB) will fuse with a lysosome to form an endolysosome, in a process promoted by the <u>ho</u>motypic fusion and <u>p</u>rotein <u>s</u>orting (HOPS) complex. Protein degradation takes place in the endolysosome (Huotari & Helenius, 2011; Spang, 2016; Szentgyörgyi & Spang, 2023).

At least the ESCRT-0 component Hrs (HGRS-1 in *C. elegans*) can already be detected on Rab5-positive endocytic vesicles (Pons *et al*, 2008; Raiborg *et al*, 2001; Solinger *et al*., 2022), indicating that cargo sorting into the degradative pathway is a very early event in the endosomal pathway. Whether and how the ESCRT machinery and Mon1/Ccz1 are coordinated remains unclear.

In this paper, we provide strong evidence for a connection between early ESCRTs, the recruitment of Mon1/Ccz1 to endosomes and Rab conversion in *C. elegans* and in mammalian cells. We show that early ESCRTs act upstream of Rab conversion and that loss of early ESCRTs prevents the recruitment of Rab7 onto endosomes. Likewise, overabundance of ubiquitinated cargoes block Rab conversion. We find that the HOPS complex can act as a Rab7GEF at least in the absence of Mon1/Ccz1 *in vivo*. We propose that the availability of ubiquitinated cargoes stabilizes the Rab5GEF Rabex5 on endosomes preventing its displacement by Mon1/Ccz1. Thus, corralling of cargoes towards sites of ILV formation by early ESCRTs is a key process in endosome maturation and a pre-requisite for Rab conversion.

## Results

### A targeted RNAi screen reveals a potential connection between Rab conversion and ESCRT function

During endosome maturation a number of processes have to happen, at least some of which should be coordinated. To test whether intraluminal vesicle (ILV) formation by the ESCRT complexes is coordinated with Rab5-to-Rab7 conversion, we performed an RNAi screen with ESCRT components in wild-type and *sand-1(KO)* mutant animals, in which Rab conversion is blocked (Poteryaev *et al*., 2010; Poteryaev *et al*., 2007; Solinger & Spang, 2014) (Fig. 1, Fig. EV1). In both strains GFP-RAB-5 and mCherry-RAB-7 are expressed in the intestine. We observed genetic interactions, such as synthetic lethality, between a subset of ESCRT components and the Rab7GEF SAND-1, suggesting a link between Rab conversion and the ESCRT machinery (Appendix Tables S2 and S3). In wild-type animals, RAB-5 is mostly localized apically, close to the gut lumen with a few puncta distributed throughout the cell, while RAB-7 is mostly on apically localized endosomes, which are bit further away from the gut lumen ((Solinger & Spang, 2014), Fig. 1A). In contrast, in *sand-1(KO)* animals, RAB-5 endosomes are enlarged and the RAB-7 localization to apical endosomes is lost (Poteryaev *et al*., 2010; Poteryaev *et al*., 2007; Solinger & Spang, 2014) (Fig. 1B) As expected, knockdown of any of the ESCRT components affected the endosomal system, and RAB-5 positive endosomes were enlarged in wild-type animals when early ESCRTs, ESCRT-0 and - I, were depleted (Fig. 1A, Fig. EV1A). Some of the knockdowns showed conspicuously similar phenotypes to the ones observed in *sand-1(KO)* (Fig. 1, Fig. EV1, Appendix Tables S4 and S5). However, *in sand-1(KO)* animals, knockdown of the early ESCRTs did not further increase the size of RAB5-positive endosomes (Fig. 1B, Fig. EV1B, Appendix Tables S4 and S5). Knockdown of late ESCRT components (ESCRT-III and ESCRT-IV) resulted in co-localization of RAB-5 and RAB-7 on endosomes in wild-type animals, indicating that Rab conversion could be initiated but not completed under these conditions. These data suggest that early ESCRTs may act upstream of Rab conversion, while late ESCRTs might act in parallel and downstream of Rab conversion. Moreover, since knock-down of early ESCRTs did not aggravate the *sand-1(KO)* mutant enlarged RAB-5-positive endosome phenotype, it is conceivable that there is coordination between Rab conversion and ILV formation.

**Figure 1:**
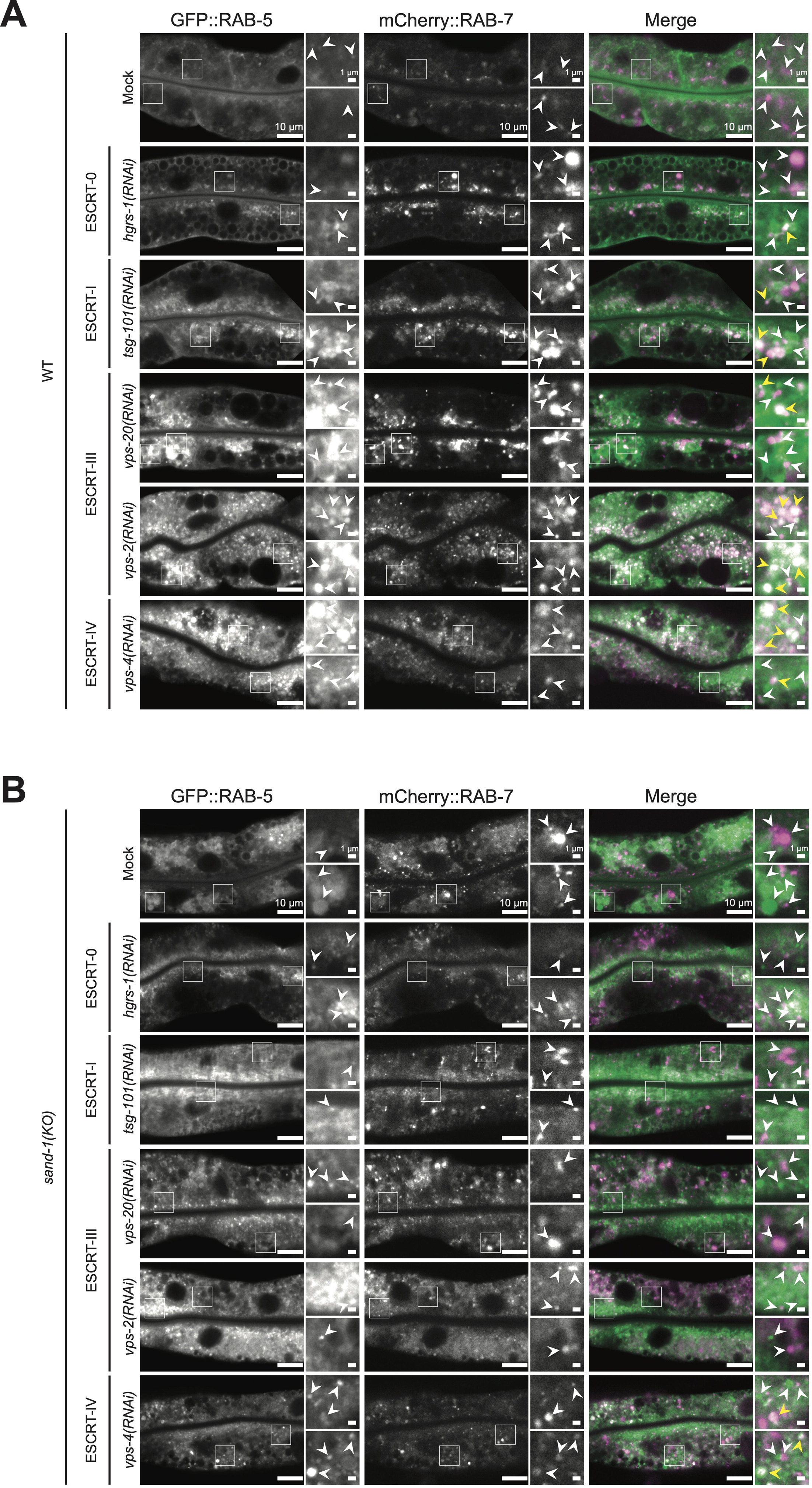
Knockdown of ESCRT mediated ILV formation impairs endosome maturation in WT and *sand-1(KO)* **A.** Knockdown of ESCRT components in a strain expressing GFP::RAB-5 and mCherry::RAB-7 in the intestine. In the merged images, colocalization events are marked via yellow arrowheads (Signals: RAB-5 > green and RAB-7 > magenta) **B.** Knockdown of ESCRT components in *sand-1(KO)* expressing GFP::RAB-5 and mCherry::RAB-7 in the intestine. White arrowheads pointing to GFP::RAB-5 and mCherry::RAB-7 positive structures, respectively in the individual channels. Colocalization events are marked via yellow arrowheads in the merges (Signals: RAB-5 > green and RAB-7 > magenta) Data information: Merged images were individually adjusted in all panels. Representative pictures with magnifications (white box) on the right are shown for each experiment (scale bars: 10 μm (main pictures) and 1 μm (insets)); *n* = 3 independent experiments.

### The number of ILVs is reduced in ESCRT knockdowns and in *sand-1(KO)* animals

To gain a better understanding of the endosomal phenotypes, we performed transmission electron microscopy (TEM) on wild-type and *sand-1(KO)* animals, in which we also knocked down an early (TSG-101) or a late (VPS-2) ESCRT component (Fig. 2, Fig. EV2 and Appendix Fig. S2). Consistent with the live cell imaging and previous studies (Frankel *et al*, 2017; Poteryaev *et al*., 2010; Poteryaev *et al*., 2007; Solinger & Spang, 2014), endosomes were enlarged in *sand-1(KO)* and in an ESCRT-I knock-down, irrespective of the strain background (Fig. EV2). In contrast ESCRT-III knockdown had a somewhat reduced effect on endosome size as also reported previously (Frankel *et al*., 2017). We defined an MVB in our micrographs as a structure that contained at least one ILV. As expected, the number of ILVs was decreased upon knock-down of ESCRT components (Fig. 2A and B). A similar effect was observed already in *sand-1(KO)* animals, but was not exacerbated by the concomitant knockdown of ESCRT components. Neither MVB nor ILV size was strongly affected in *sand-1(KO)* animals (Fig. 2C-F), consistent with the notion that the enlarged endosomes in *sand-1(KO)* animals are RAB-5 positive (Poteryaev *et al*., 2010; Poteryaev *et al*., 2007). Taken together, our data so far are consistent with a possible coordination between Rab conversion and ILV formation.

**Figure 2:**
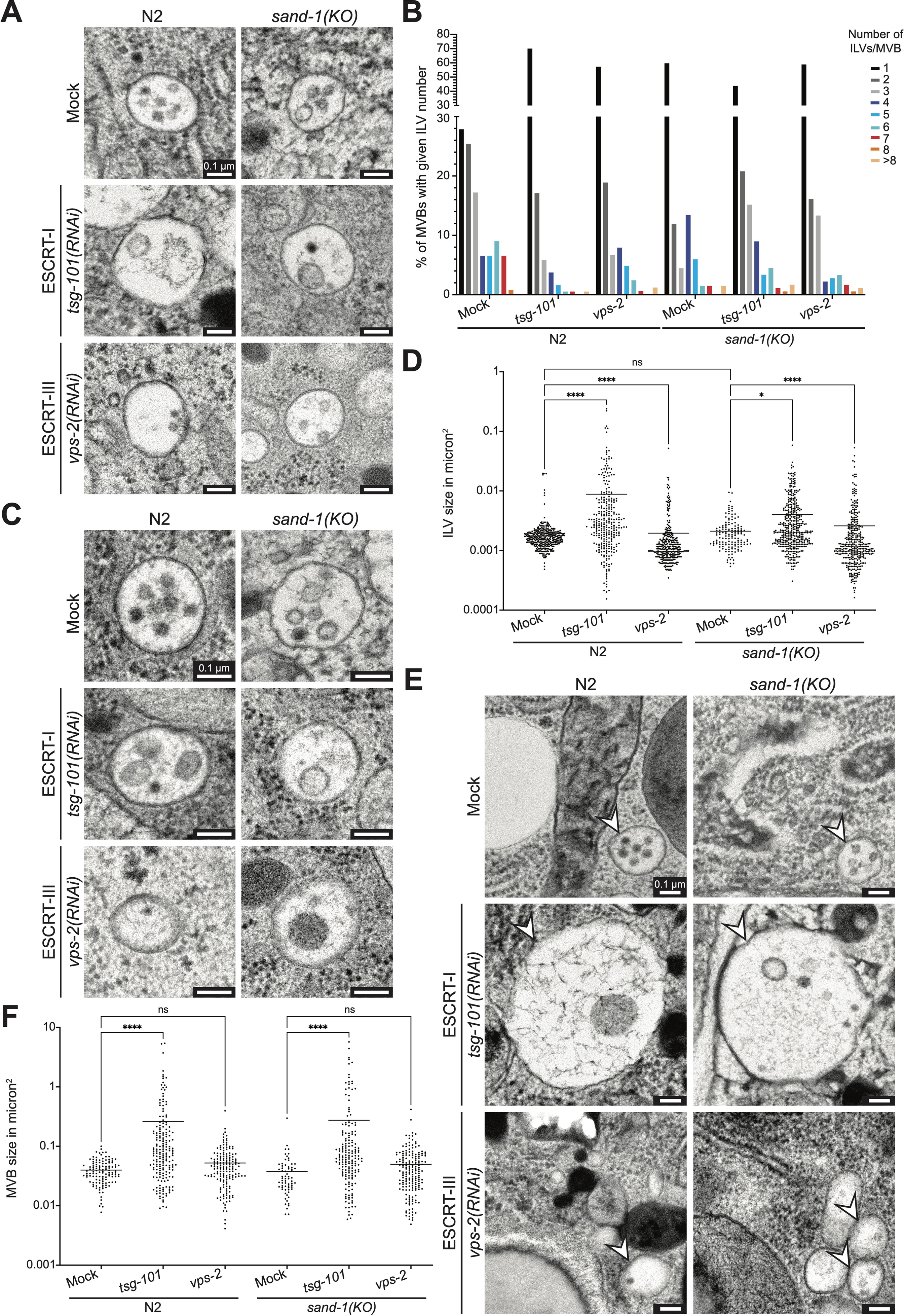
ESCRT knockdowns affect the MVB and ILV size but not the ILV number in WT. **A.** Knockdowns of *tsg-101* or *vps-2* reduce the number of ILVs in WT animals. *sand-1(KO)* reduces ILV number independently of ESCRT knockdown. **B.** Quantification of the ILV phenotypes shown in (**A**). ILVs/MVB were counted in three worms per condition (*n* = 3) and summed in 9 categories, 1-8 ILVs/MVB and more than 8 ILVs/MVB. The results per category are shown in % of the total amount as bar graph. **C.** *tsg-101(RNAi)* causes in both strains an increase in ILV size. The *vps-2(RNAi)* resulted in the formation of smaller and bigger structures altering the ILV size range in both strains. *sand-1* knockout alone does not change the ILV size. **D.** Quantification of the ILV size phenotypes depicted in (**C**). The size of each ILV was measured in μm^2^ in three worm intestines per condition (*n* = 3). Data are shown in log 10 scale (Y-axis) with corresponding mean value as scatter plot. *P*-values: Mock vs. *tsg-101(RNAi) P* < 0.0001, Mock vs. *vps-2(RNAi) P* < 0.0001, Mock vs. *sand-1(KO) P* > 0.9999, *sand-1(KO)* vs. *sand-1(KO)* + *tsg-101(RNAi) P* = 0.0439, *sand-1(KO)* vs. *sand-1(KO)* + *vps-2(RNAi) P* < 0.0001. **E.** MVB size phenotypes in WT and *sand-1(KO)* caused by *tsg-101(RNAi)* or *vps-2(RNAi)*. Knockdown of *tsg-101* but not of *vps-2* causes the formation of increased MVBs in both strains. The *sand-1* knockout alone does not change the MVB size. Representative MVBs are highlighted with white arrowheads in each picture. **F.** The MVB size was quantified via measurement of the MVB area in μm^2^ in three worm intestines per condition (*n* = 3). Displayed quantification belongs to (**E**). Data are shown in log 10 scale (Y-axis) with corresponding mean values as scatter plot. *P*-values: Mock vs. *tsg-101(RNAi) P* < 0.0001, Mock vs. *vps-2(RNAi) P* > 0.9999, Mock vs. *sand-1(KO) P* > 0.9999, *sand-1(KO)* vs. *sand-1(KO)* + *tsg-101(RNAi) P* < 0.0001, *sand-1(KO)* vs. *sand-1(KO)* + *vps-2(RNAi) P* = 0.4844. Data information: Pictures are individually adjusted (scale bars: 0.1 μm). Representative MVBs are shown for each examined condition (**A**, **C** and **E**). Kruskal-Wallis tests with multiple comparisons were performed to determine the statistical significance between the examined conditions (**D** and **F**). Significance levels are displayed in the graphs as * *P* ≤ 0.05, **** *P* ≤ 0.0001 and ns *P* > 0.05. Unprocessed images and statistical raw data are available as source data.

### HGRS-1 is present on RAB-5 positive and absent on RAB-7 positive endosomes

Our screen indicated that early ESCRTs act upstream of Rab conversion. Moreover, in mammalian cells the residence time of ESCRT-0 HRS (HGRS-1 in *C. elegans*) on endosomes is prolonged, when ESCRT-III components are depleted (Quinney *et al*, 2019). However, it remains unclear whether the HRS-positive endosomes are Rab5 or Rab7 positive under these conditions. To address this issue, we knocked-down ESCRT-I, -III and -IV components in worms expressing GFP-HGRS-1 together with either RFP-RAB-5 or mCherry-RAB-7. HGRS-1 co-localized with RAB-5, even when late ESCRT components were depleted (Fig. 3A-D), while we did not observe co-localization between HGRS-1 and RAB-7 (Fig. 3E-G). Thus, the presence of HGRS-1 and RAB-7 on endosomes appears to be mutually exclusive, suggesting that HGRS-1 might have to leave endosomes prior to Rab conversion.

**Figure 3:**
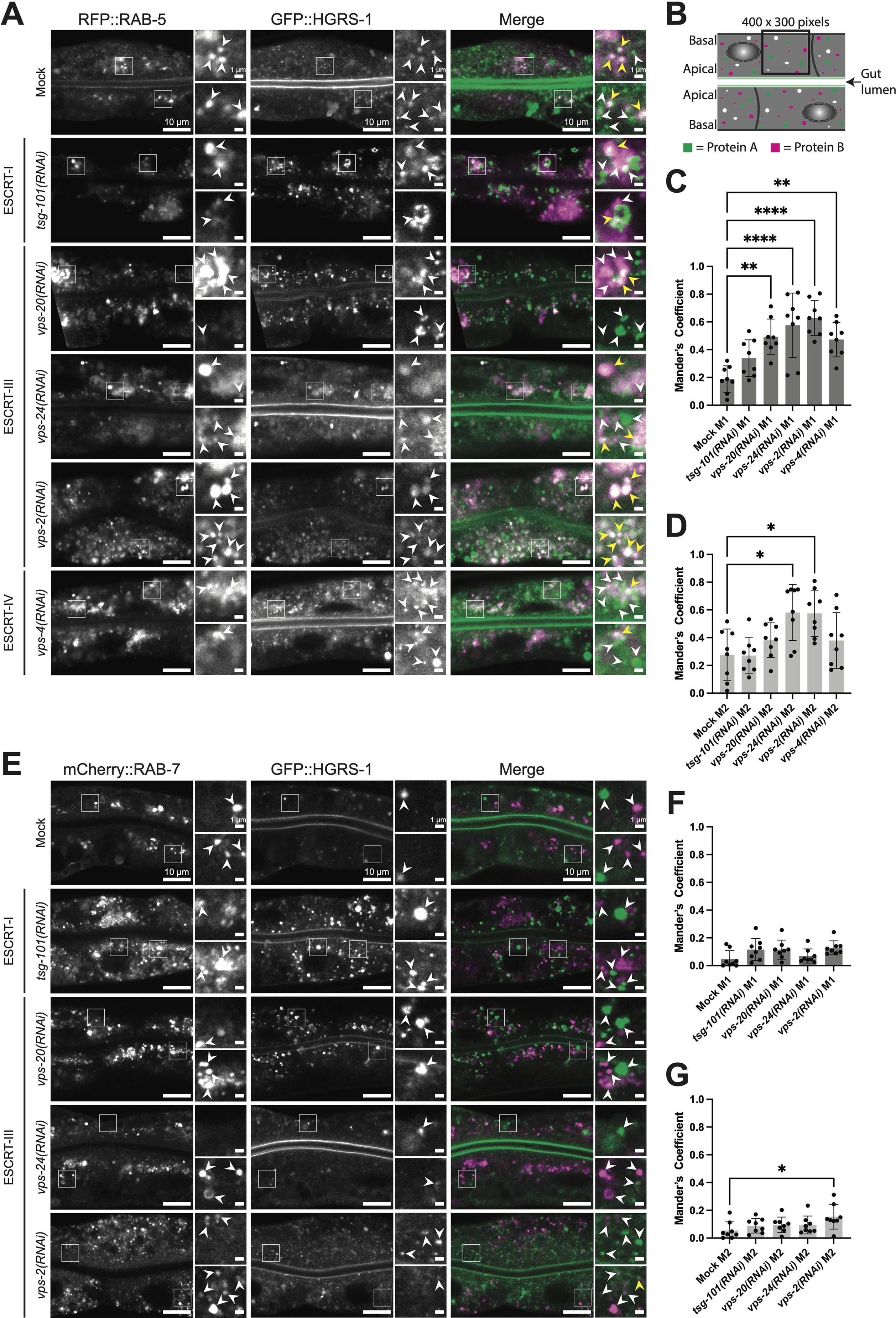
HGRS-1 and RAB-5 but not RAB-7 show high colocalization on endosomes if ILV formation is impaired. **A.** GFP::HGRS-1 colocalizes with RFP::RAB-5 under early and late ESCRT factor knockdowns and is predominantly located on enlarged structures under *tsg-101, vps-20, vps-2* and *vps-4* conditions. White arrowheads marking GFP::HGRS-1 and RFP::RAB-5 positive structures, respectively in the individual channels. Yellow arrowheads marking colocalization events in the merges (Signals: HGRS-1 > green and RAB-5 > magenta). **B.** Schematical representation of a *C. elegans* gut with polarized intestinal cells and two expressed marked example proteins (Protein A in green and Protein B in magenta) which show partial colocalization (white). The colocalization quantification approach using a ROI (400 x 300 pixels) used in **C**, **D**, **F** and **G** is shown as an example. **C**, **D**. Colocalization quantification of GFP::HGRS-1 and RFP::RAB-5 belonging to (**A**). For analysis shown in (**C** and **D**), 8 worms (*n* = 8) were examined per condition (*n* = 3 independent experiments). Analysis of colocalization via Mander’s coefficient (M1: RFP::RAB-5 signal overlapping with GFP::HGRS-1 signal, M2: GFP::HGRS-1 signal overlapping with RFP::RAB-5 signal).(**C**) Obtained *P-*values: Mock M1 vs. *tsg-101(RNAi)* M1 *P* = 0.3273, Mock M1 vs. *vps-20(RNAi)* M1 *P* = 0.002, Mock M1 vs. *vps-24(RNAi)* M1 *P* = < 0.0001, Mock M1 vs. *vps-2(RNAi)* M1 *P* < 0.0001, Mock M1 vs. *vps-4(RNAi)* M1 *P* = 0.0044; (**D**) Obtained *P-*values: Mock M2 vs. *tsg-101(RNAi)* M2 *P* > 0.9999, Mock M2 vs. *vps-20(RNAi)* M2 *P* = 0.8217, Mock M2 vs. *vps-24(RNAi)* M2 *P* = 0.0114, Mock M2 vs. *vps-2(RNAi)* M2 *P* = 0.0139, Mock M2 vs. *vps-4(RNAi)* M2 *P* = 0.8373. **E**. The very minor colocalization of GFP::HGRS-1 and mCherry::RAB-7 is only slightly affected under early or late ESCRT knockdown conditions and never reaches the levels of GFP::HGRS-1 and RFP::RAB-5 shown in (**A**). In the close ups white arrowheads marking GFP::HGRS-1 and mCherry::RAB-7 positive structures, respectively in the individual channels. Merges showing a combination of both channels (Signals: HGRS-1 > green and RAB-7 > magenta). Colocalization is indicated via yellow arrowhead. **F**, **G**. Colocalization quantification of GFP::HGRS-1 and mCherry::RAB-7 belonging to (**E**). For analysis shown in (**F** and **G**), 8 worms (*n* = 8) were examined per condition (*n* = 3 independent experiments). Analysis of colocalization via Mander’s coefficient (M1: mCherry::RAB-7 signal overlapping with GFP::HGRS-1 signal, M2: GFP::HGRS-1 signal overlapping with mCherry::RAB-7 signal).(**F**) Obtained *P-*values: Mock M1 vs. *tsg-101(RNAi)* M1 *P* = 0.38, Mock M1 vs. *vps-20(RNAi)* M1 *P* = 0.3421, Mock M1 vs. *vps-24(RNAi)* M1 *P* > 0.9999, Mock M1 vs. *vps-2(RNAi)* M1 *P* = 0.0907; (**G**) Obtained *P-*values: Mock M2 vs. *tsg-101(RNAi)* M2 *P* > 0.9999, Mock M2 vs. *vps-20(RNAi)* M2 *P* > 0.9999, Mock M2 vs. *vps-24(RNAi)* M2 *P* > 0.9999, Mock M2 vs. *vps-2(RNAi)* M2 *P* = 0.0247. Data information: Merges were individually adjusted in all panels. Representative pictures with Enlargements (white box) on the right are shown for each experiment (scale bars: 10 μm (main pictures) and 1 μm (close ups)). Ordinary one-way ANOVA with Tukey’s multiple comparisons test was performed for each experiment to determine the statistical significance between the examined conditions (**C** and **D**). For (**F** and **G**) a Kruskal-Wallis test with Dunn’s multiple comparisons test was performed for each experiment for the same purpose. Significance levels are displayed in the graphs as * *P* ≤ 0.05, ** *P* ≤ 0.01, *** *P* ≤ 0.001, **** *P* ≤ 0.0001 and ns *P* > 0.05. Each data point in (**C** and **D**) and in (**F** and **G**) represents the colocalization of the markers measured in one worm. In the graphs (**C**, **D**, **F** and **G**) the mean ± s.d. is shown for all conditions. Unprocessed images and statistical raw data are available as source data.

### The level of ubiquitinated cargo on endosomes impacts Rab conversion

The major role of ESCRT-0 is the corralling of ubiquitinated cargoes on early endosomes and hand them over to downstream ESCRTs for inclusion into ILVs. HRS binds directly to ubiquitinated cargo, and its presence on endosomes is strongly influenced by this function (Hirano *et al*, 2006; Komada *et al*, 1997; Raiborg *et al*, 2002; Wollert & Hurley, 2010). Accumulation of ubiquitinated cargoes would lead to persistent HRS localization on endosomes and thereby hamper endosomes maturation. To test this possibility, we first assessed whether ubiquitinated cargoes accumulate on endosomes when ESCRT levels are reduced by using a *C. elegans* strain expressing ubiquitin fused to GFP (GFP-UBQ) (Bakowski *et al*, 2014). Rather than looking at an individual cargo, we decided to look at ubiquitin on endosomes to more globally assess cargo effects. In control animals, GFP-UBQ is present in a haze with some occasional foci, presumably endosomes, in the cytoplasm and concentrated in the nucleus (Fig. 4A, Fig. EV3A). Knockdown of ESCRTs locked ubiquitin on endosomes and the nuclear pool was depleted (Fig. 4A and B, Fig. EV3A). Of note, depleting HGRS-1 or VPS-20 (ESCRT-III; CHMP-6 in mammals) led to a reduction of the GFP-UBQ signal (Fig. EV3B), suggesting that some ubiquitin could be degraded under these conditions. Moreover, the size of the GFP-UBQ positive endosomes was enlarged (Fig. EV3C). We also aimed to increase ubiquitinated cargo on endosomes by depleting the deubiquitinase USP-50, which removes ubiquitin from cargoes before they enter ILVs (Clague & Urbé, 2006; Henne *et al*., 2011). However, unfortunately, HGRS-1 is itself ubiquitinated (Katz *et al*, 2002; Polo *et al*, 2002; Stringer & Piper, 2011) and is also a substrate of USP-50 (Row *et al*, 2006; Zhang *et al*, 2014). Hence upon *usp-50(RNAi),* HGRS-1 remains ubiquitinated and is degraded (Fig. EV3D). Still, our data suggest that reduction of ubiquitinated cargoes on endosomes may allow Rab conversion to occur. To test this hypothesis, we reduced the ubiquitin levels by RNAi in the *sand-1(KO)* mutant. We observed a decrease in size of the RAB-5 positive endosomes and RAB-7 was partially recruited to apically localized endosomes (Fig. 5A-C, Fig. EV4C). RAB-7 recruitment onto endosomes was sufficient to release the cargo hTFR-GFP from being trapped in internal vesicles (Fig. 5D-H) and also improved the morphology of the LMP-1 positive compartments (Fig. 5I-K, Fig. EV4B and C). Moreover, the colocalization of LMP-1 and RAB-7 remained high (Fig. 5L). Consistent with these findings, knockdown of *usp-50* in wild-type animals let to mislocalization of RAB-5 and co-localization of RAB-5 with RAB-7 and GFP-UBQ, respectively (Fig. EV5A-D). Furthermore, knockdowns of *hgrs-1* and *usp-50* showed negative genetic interaction with *sand-1(KO)* (Appendix Table S2, S3, S6 and S7). Our data suggest that in the absence of ESCRT-0, or when ubiquitinated cargo cannot be corralled and accumulates on endosomes, endosomal flow is abrogated and Rab conversion is blocked. Moreover, relief from the accumulation of ubiquitinated cargo seemed to allow Rab conversion, even in the absence of the Rab7GEF SAND-1.

**Figure 4:**
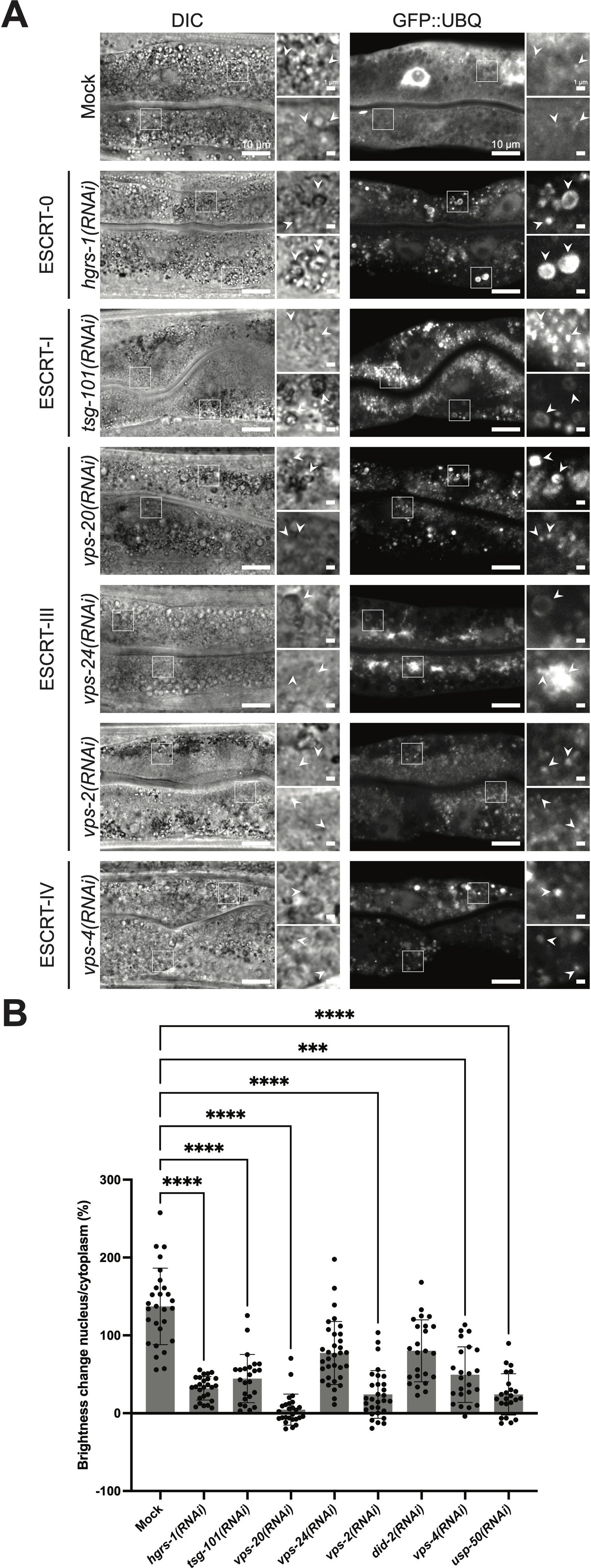
Early and late ESCRT knockdowns affect UBQ localization in intestinal cells. **A.** Knockdowns of core early ESCRT factors (0 and I) and late ESCRT factors (III-IV) cause a reduction in GFP::UBQ levels in the nucleus to varying degrees. White arrowheads indicating GFP::UBQ positive structures and the belonging region in the DIC. **B.** Quantification of the nuclear GFP::UBQ amount with respect to the cytoplasmic signal in (**A**) and Fig. EV3A. Ten worms (*n* = 10) were quantified (*n* = 3 independent experiments). Each data point represents the result of one nucleus/ cytoplasm comparison in the indicated condition. *P-*values: Mock vs. *hgrs-1(RNAi) P* < 0.0001, Mock vs. *tsg-101(RNAi) P* < 0.0001, Mock vs. *vps-20(RNAi) P* < 0.0001, Mock vs. *vps-24(RNAi) P* = 0.1914, Mock vs. *vps-2(RNAi) P* < 0.0001, Mock vs. *did-2(RNAi) P* = 0.5219, Mock vs. *vps-4(RNAi) P* = 0.000113, Mock vs. *usp-50(RNAi) P* < 0.0001. Data information: Shown are representative pictures with magnifications (white box) on the right for each experiment (scale bars: 10 μm (main pictures) and 1 μm (inlet)). Corresponding DIC pictures elucidate the extent of the gut, marker independent. GFP channel pictures of *hgrs-1* and *vps-20* knockdowns are increased in brightness (for analogous pictures with equal brightens see Fig. EV3). Kruskal-Wallis test with Dunn’s multiple comparisons test was performed for each experiment to determine the statistical significance between the examined conditions (**B**). Significance levels are displayed in the graphs as * *P* ≤ 0.05, ** *P* ≤ 0.01, *** *P* ≤ 0.001, **** *P* ≤ 0.0001 and ns *P* > 0.05. The data show the mean ± s.d. for all examined conditions. Unprocessed images and statistical raw data are available as source data.

**Figure 5:**
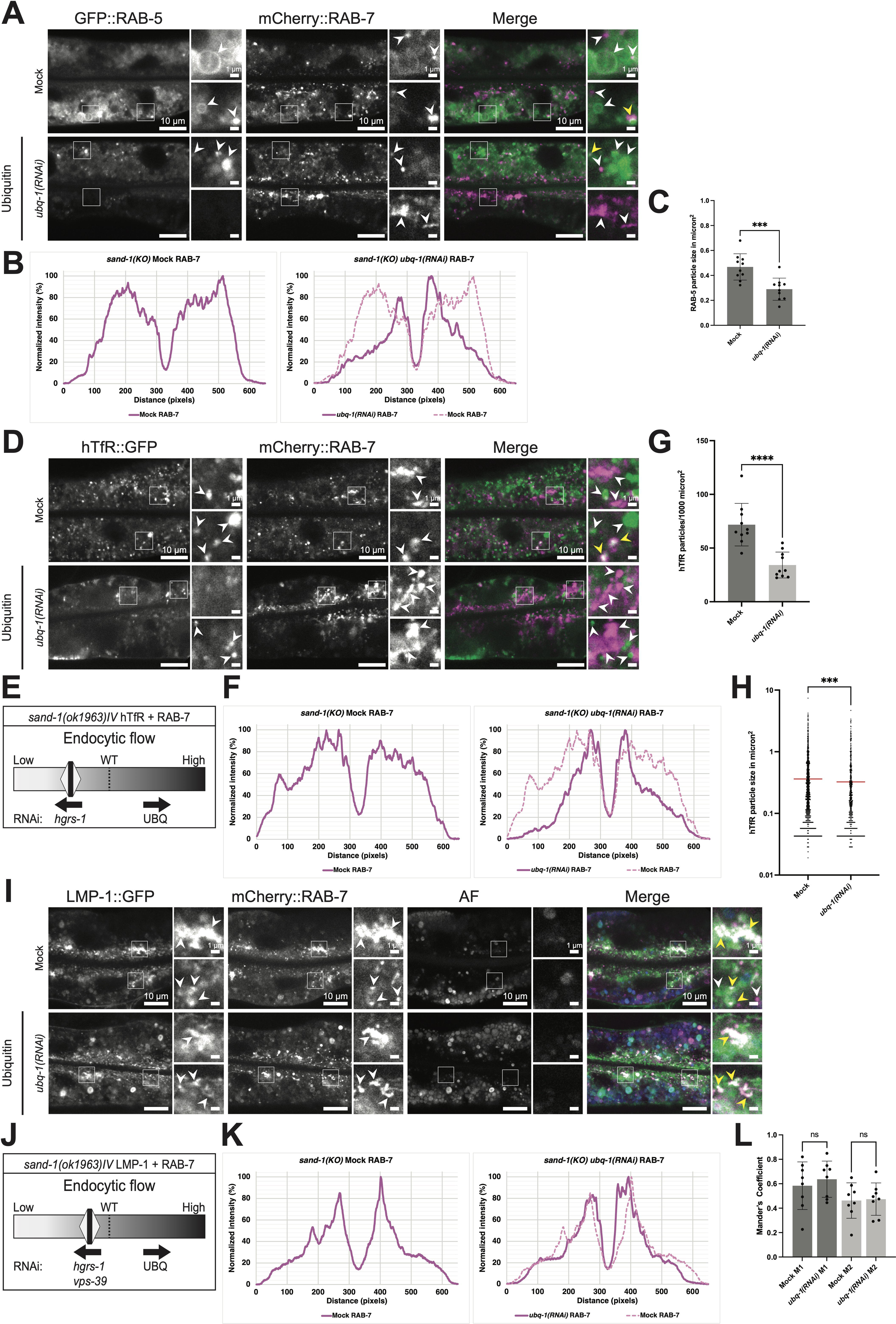
The reduction of UBQ levels rescues RAB-7 endosome association *sand-1(KO)*. **A.** The reduction of UBQ levels in *C elegans* intestinal cells causes an accumulation of mCherry::RAB-7 on the apical side and reduces the size of GFP::RAB-5 positive structures in *sand-1(KO)*. White arrowheads pointing to GFP::RAB-5 and mCherry::RAB-7 positive structures, respectively in the individual channels. Co-localization events are highlighted with yellow arrowheads in the merges (Signals: RAB-5 > green and RAB-7 > magenta). **B**, **C**. Quantification of the mCherry::RAB-7 area localization (**B**) and GFP::RAB-5 particle size in micron^2^ (**C**) belonging to (**A**). For analyses shown in (**B**) 5 (*n* = 5), respectively 10 (*n* = 10) (**C**) worms were examined per condition (*n* = 3 independent experiments). In (**C**) each data point represents the mean particle size of one measured worm. *P-*value for (**C**): Mock vs. *ubq-1*+Control*(RNAi) P* = 0.000103. **D.** Reduction of UBQ levels in a *sand-1(KO)* mutant overexpressing an artificial endocytic cargo rescues mCherry::RAB-7 localization. The hTfR::GFP signal is shown with increased brightness in case of UBQ reduction condition to compensate for the reduced signal (for same brightness see Appendix Fig. S3). White arrowheads pointing to hTfR::GFP:: and mCherry::RAB-7 positive structures, respectively in the individual channels. Merges show colocalization under Mock conditions with yellow arrowheads (Signals: hTfR > green and RAB-7 > magenta). **E.** Schematic representation of the impaired endocytic flow in the *sand-1(KO)* overexpressing tagged hTfR and RAB-7. Effects of UBQ reduction and *hgrs-1(RNAi)* are indicated with black arrows. **F-H**. mCherry::RAB-7 area localization quantification (**F**) and quantification of the hTfR::GFP particle abundance (per 1000 micron^2^) (**G**) and size (in micron^2^) (**H**) shown in (**D**). For analyses shown in (**F**) 5 (*n* = 5), (**G**) 10 (*n* = 10) and (**H**) 10 (*n* = 10) worms were examined per condition (*n* = 3 independent experiments). In (**G**) each data point represents the particle abundance of one examined worm and in (**H**) all measured particles of the 10 worms per condition are shown (*n* > 2500). (**G**) *P*-values: Mock vs. *ubq-1(RNAi) P* < 0.0001; (**H**) *P*-values: Mock vs. *ubq-1(RNAi) P* = 0.000110. **I.** Reduction of UBQ levels, partially rescues the mCherry::RAB-7 localization in *sand-1(KO)*. White arrowheads point to LMP-1::GFP and mCherry::RAB-7 positive structures, respectively in the individual channels. The AF channel depicts autofluorescent gut granules, also visible in the GFP channel and the merge. In the merges colocalization is shown with yellow arrowheads (Signals: LMP-1 > green, RAB-7 > magenta and AF > blue). **J.** Schematic representation of the endocytic flow in the *sand-1(KO)* strain overexpressing tagged LMP-1 and RAB-7. The consequences of the UBQ reduction and *hgrs-1(RNAi)* are shown as black arrows. **K, L**. Quantification of the mCherry::RAB-7 area localization (**K**) (equivalent quantification of LMP-1::GFP is shown in Fig. EV4B and C) and LMP-1::GFP/mCherry::RAB-7 colocalization (**L**) shown in (**I**). For analyses shown in (**K**) 5 (*n* = 5), respectively 8 (*n* = 8) (**L**) worms were examined per condition (*n* = 3 independent experiments). (**L**) each data point represents the co-localization of the markers measured in one worm. Analysis of co-localization via Mander’s coefficient (M1 mCherry::RAB-7 signal overlapping with LMP-1::GFP signal, M2: LMP-1::GFP signal overlapping with mCherry::RAB-7 signal). (**L**) *P-*values: Mock M1 vs. *ubq-1(RNAi)* M1 *P* = 0.9923, Mock M2 vs. *ubq-1(RNAi)* M2 *P* > 0.9999. Data information: Merged images were individually adjusted in all panels. Representative pictures with magnifications (white box) on the right are shown for each experiment (scale bars: 10 μm (main pictures) and 1 μm (inset)). Ordinary one-way ANOVA with Tukey’s multiple comparisons test was performed for each experiment to determine the statistical significance between the examined conditions (**C** and **L**). For (**G** and **H**) a Mann-Whitney test (two-tailed) was performed for each experiment for the same purpose. Significance levels are displayed in the graphs as *** *P* ≤ 0.001, **** *P* ≤ 0.0001 and ns *P* > 0.05. The data show the mean ± s.d. for all examined conditions (**C**, **G** and **L**). In (**H**) the mean is shown for each condition as red line in the graph. Unprocessed images and statistical raw data are available as source data.

### HOPS is a Rab7GEF

We showed above that reducing ubiquitination on endosomal cargo partially rescues the block in Rab conversion in the *sand-1(KO)* mutant. The SAND-1/CCZ-1 complex (Mon1/Ccz1 in mammals and yeast) is the GEF for RAB-7 (Nordmann *et al*, 2010). Therefore, another Rab7GEF must exist. Previously, the HOPS complex component Vps39 was proposed to act as an Rab7GEF (Binda *et al*, 2009; Wurmser *et al*, 2000), and HOPS activated yeast Rab7 *in vitro*, albeit less efficiently than Mon1/Ccz1 (Nordmann *et al*., 2010), but still this function remained controversial. We tested whether VPS-39 acts as a Rab7GEF in our system. Therefore, we repeated the ubiquitin depletion in *sand-1(KO)*, but this time, we co-depleted VPS-39. Under these conditions, RAB-7 recruitment on apical endosomes was strongly reduced (Fig. 5A and B, Fig. 6A and B, Fig. EV4F). These results indicate that VPS-39 is indeed a Rab7GEF, and can activate RAB-7 on endosomes when the ubiquitinated cargo load is low. The defect in endosome maturation in *sand-1(KO)* can be partially rescued by overexpression of RAB-7, as indicated by the LMP-1 localization (Fig. 5I and K, Fig. EV4B). Depletion of VPS-39 reversed this partial rescue and as a consequence, RAB-7 was dispersed and LMP-1 accumulated again (Fig. 6C and D, Fig. EV4D). Moreover, the colocalization of RAB-7 and LMP-1 was significantly reduced in *vps-39*(RNAi) animals indicating that VPS-39 is required for efficient recruitment of RAB-7 to LMP-1 positive structures in *sand-1(KO)* (Fig. 6E). Finally, *vps-39* displayed a negative genetic interaction with *sand-1(KO)*, a phenotype which was not rescued by concomitant depletion of ubiquitin or overexpression of RAB-7 (Appendix Tables S6 and S7). Taken together, our data indicate that the HOPS component VPS-39 can act as Rab7GEF in the *C. elegans* intestine and that this activity is essential for the viability of *sand-1(KO)* animals.

**Figure 6:**
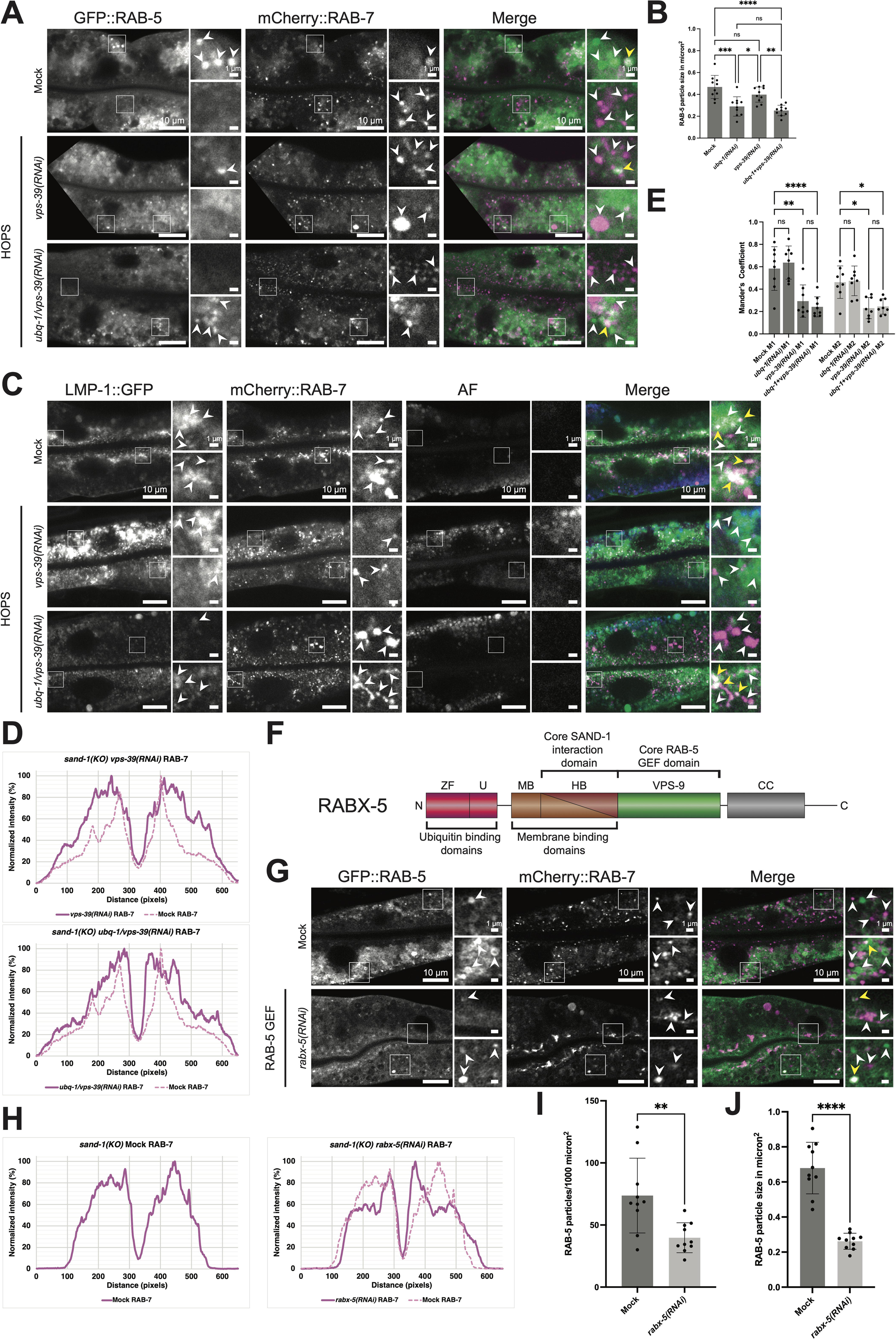
RABX-5 knockdown rescues *sand-1(KO)* and VPS-39 acts as second RAB-7 GEF *in vivo*. **A.** The knockdown of *vps-39* abrogates the rescue of UBQ reduction in *sand-1(KO)* animals. White arrowheads point to GFP::RAB-5 and mCherry::RAB-7 positive structures, respectively in the individual channels. Colocalization events are marked with yellow arrowheads in the merges (Signals: RAB-5 > green and RAB-7 > magenta). **B.** Quantification of the GFP::RAB-5 particle size in micron^2^ belonging to (A). Control group and *ubq-1* knockdown are already shown in Fig. 5C and experimental details are the same like explained in the figure legend belonging to Fig. 5C. *P*-values: Mock vs. *vps-39(RNAi) P* = 0.2302, Mock vs. *ubq-1*+*vps-39(RNAi) P* < 0.0001, *ubq-1*+Control*(RNAi)* vs. *vps-39(RNAi) P* = 0.0240, *ubq-1*+Control*(RNAi)* vs. *ubq-1*+*vps-39(RNAi) P* = 0.7328, *vps-39(RNAi)* vs. *ubq-1*+*vps-39(RNAi) P* = 0.0015. **C.** In *vps-39* knockdown cells, LMP-1::GFP forms accumulations and mCherry::RAB-7 shows a dispersed pattern. These patterns are only partial rescuable via UBQ reduction in this *sand-1(KO)*. White arrowheads pointing to LMP-1::GFP and mCherry::RAB-7 positive structures, respectively in the individual channels. In addition, the AF channel depicts the gut granules also visible in the GFP channel. Colocalization events are marked with yellow arrowheads in the merges (Signals: LMP-1 > green, AF > blue and RAB-7 > magenta). **D.** Quantification of the mCherry::RAB-7 area localization (equivalent quantification of LMP-1::GFP is shown in Fig. EV4D and E) belonging to (**C**). The control group is the same as shown in Fig. 5L and the data were collected and analyzed like described there. **E.** Colocalization quantification of LMP-1::GFP and mCherry::RAB-7 belonging to (C). Control group and *ubq-1* knockdown as displayed in Fig. 5L and experimental details are visible in the figure legend Fig. 5L. *P*-values: Mock M1 vs. *vps-39(RNAi)* M1 *P* = 0.0013, Mock M1 vs. *ubq-1*+*vps-39(RNAi)* M1 *P* < 0.0001, *vps-39(RNAi)* M1 vs. *ubq-1*+*vps-39(RNAi)* M1 *P* = 0.9941, Mock M2 vs. *vps-39(RNAi)* M2 *P* = 0.0198, Mock M2 vs. *ubq-1*+*vps-39(RNAi)* M2 *P* = 0.0377, *vps-39(RNAi)* M2 vs. *ubq-1*+*vps-39(RNAi)* M2 *P* > 0.9999. **F.** Graphical illustration of *C. elegans* RABX-5. RABX-5 harbors following domains from N- to C-terminus required for its function: UBQ binding > zinc finger motif (ZF) and UBQ interacting motif (U) (slight cardinal red), Membrane binding > membrane binding domain (MB) (slight brown) and helical bundle (HB) (dark brown), Core SAND-1 interaction > HB, Core RAB-5 GEF > VPS-9 (dark green), Autoinhibition > coiled coil (CC) domain (dark grey). **G.** The knockdown of *rabx-5* cause an accumulation of GFP::RAB-5 and mCherry:: RAB-7 in the apical area of the worm gut and a reduction in size and abundance of GFP::RAB-5 structures in *sand-1(KO)*. White arrowheads pointing to GFP::RAB-5 and mCherry::RAB-7 positive structures, respectively in the individual channels. Colocalization events are marked with yellow arrowheads in the merges (Signals: RAB-5 > green and RAB-7 > magenta). **H-J**. Quantification of the mCherry::RAB-7 area localization (**H**), GFP::RAB-5 particle abundance (per 1000 micron^2^) (**I**) and size (in micron^2^) (**J**) belonging to (**F**). For analysis shown in (**G**) 5 (*n* = 5), (**I**) 10 (*n* = 10) and (**J**) (*n* = 10) worms were examined per condition (*n* = 3 independent experiments). In (**I**) each data point represents the particle abundance of one examined worm and in (**J**) each data point shows the mean particle size of one measured worm. (**I**) *P*-value: Mock vs. *rabx-5(RNAi) P* = 0.0063; (**J**) *P*-value: Mock vs. *rabx-5(RNAi) P* < 0.0001. Data information: Merges were individually adjusted in all panels. Representative pictures with magnification (white box) on the right are shown for each experiment (scale bars: 10 μm (main pictures) and 1 μm (insets)). A detailed description of the performed statistics for (**B** and **E**) is available in the figure legend of Fig. 5. Welch’s t test (two-tailed) was performed for each experiment to determine the statistical significance between the examined conditions (**I** and **J**). Significance levels are displayed in the graphs as * *P* ≤ 0.05, ** *P* ≤ 0.01, *** *P* ≤ 0.001, **** *P* ≤ 0.0001 and ns *P* > 0.05. The data show the mean ± s.d. for all examined conditions (**C**, **E**, **I** and **J**). Unprocessed images and statistical raw data are available as source data.

### RABX-5 prevents Rab conversion in the presence of ubiquitinated cargo

Next, we wanted to determine the link between the ubiquitinated cargoes, early ESCRTs and Rab conversion. Besides ESCRT-0, the GEF for RAB-5, RABX-5 (Rabex5 in mammals) has ubiquitin binding properties (Fig. 6F)(Dwivedi *et al*, 2011; Lee *et al*, 2006; Mattera *et al*, 2006; Penengo *et al*, 2006). In fact, together with the membrane-binding domain, the ubiquitin binding domain is required for the Rabex5 localization on early endosomes (Lauer *et al*, 2019; Mattera & Bonifacino, 2008; Mattera *et al*., 2006; Zhu *et al*, 2007). We have shown previously that SAND-1 interacts with and displaces RABX-5 from endosomes (Poteryaev *et al*., 2010). Thus, RABX-5 would be an ideal candidate to link the reduction of ubiquitinated cargo to Rab conversion. We reasoned that as long as RABX-5 would be able to bind to ubiquitin, SAND-1 would be unable to displace it and hence, Rab conversion would be blocked. Once HGRS-1, together with other early ESCRTs, has corralled all ubiquitinated cargo into a domain for ILV formation, RABX-5 would no longer bind to ubiquitin and hence SAND-1 would be able to displace RABX-5 and initiate Rab conversion. To test this hypothesis, we knocked down RABX-5 in *sand-1(KO)* animals. Similar to the knock-down of ubiquitin (Fig. 5), the RAB-7 localization to apical endosomes was restored (Fig. 6G and H). Moreover, the size and number of the RAB-5 positive endosomes was reduced (Fig. 6G, I and J). Thus, our data indicate that RABX-5 is required for the coordination between early ESCRT function and Rab conversion by sensing the uncorralled ubiquitinated cargo on endosomes.

### The coordination between cargo corralling by ESCRT-0 and Rab conversion is conserved in mammalian cells

Finally, we wanted to test whether the crosstalk between early ESCRTs and Rab conversion is conserved in metazoans. First, we established that CRISPR-Cas9 knockout lines for ESCRT-0 (HRS KO) and ESCRT-III (CHMP6 KO) led to the enlargement of Rab5 positive endosomes, like in the knockdown of the corresponding proteins in *C. elegans* (Fig. 7A-C, Fig. 1A, Fig. 3A). Likewise, we confirmed that KO of the Rab7GEF component CCZ1 displayed enlarged Rab5 positive endosomes (Fig. 7D and F) (Podinovskaia *et al*., 2021). Interestingly, similar to *C. elegans*, over expression of Rab7 rescued the enlarged Rab5 positive endosomes phenotype in CCZ1 KO cells (Fig. 7E-G, Fig. 5I), indicating that besides MON1/CCZ1 another Rab7GEF must exist also in mammalian cells, most likely Vps39. Finally, we overexpressed HRS in CCZ1 KO cells, which rescued the enlarged Rab5-positive endosome phenotype (Fig. 7H-J). Taken together our data provide strong evidence for an evolutionary conserved mechanism by which ESCRT-0 function is required upstream of Rab conversion and that together with ubiquitinated cargo levels ESCRT-0 may serve as a timer for the initiation of Rab conversion.

**Figure 7:**
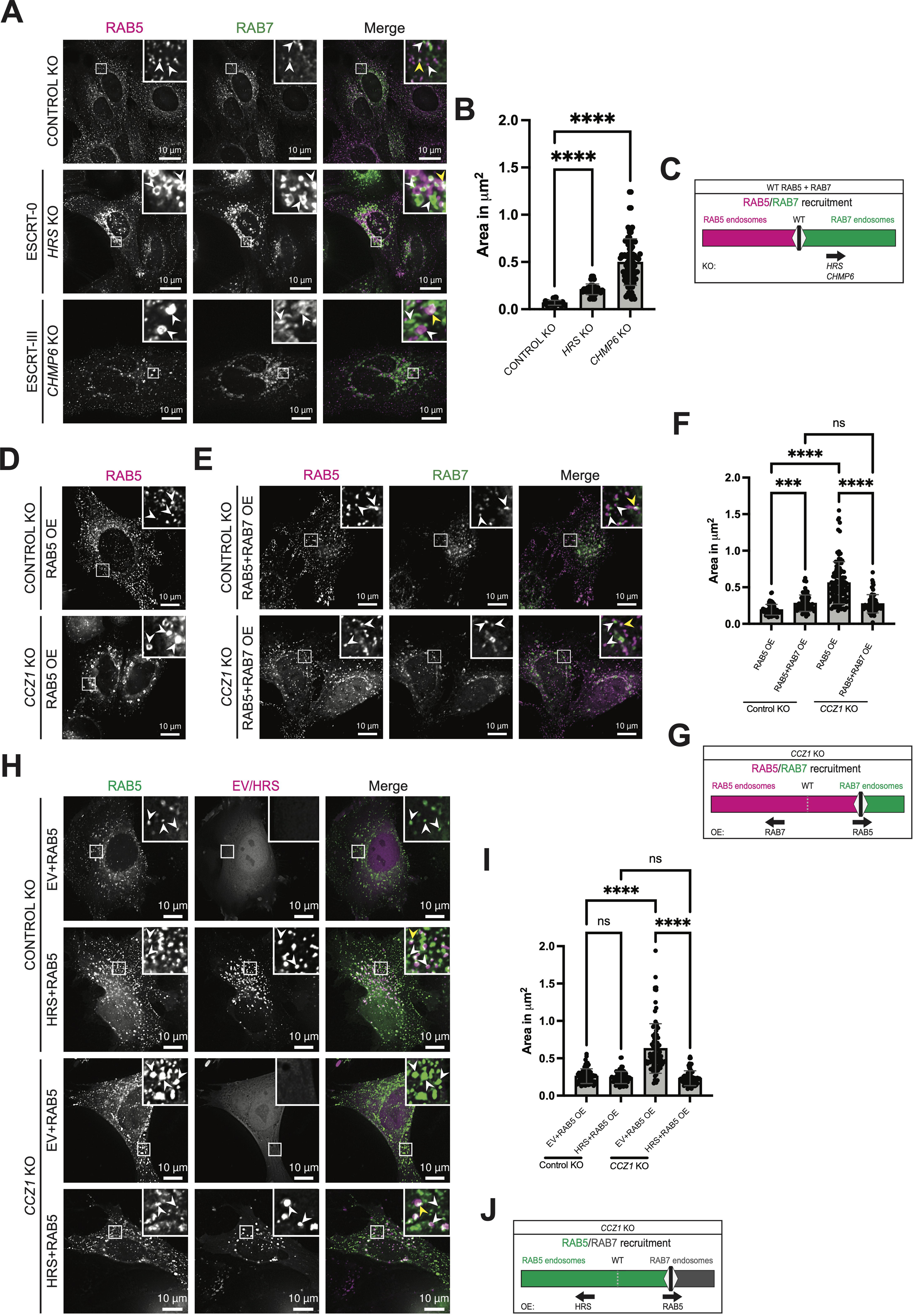
The coordination of ESCRT mediated ILV formation and Rab conversion and the second Rab7GEF downstream of MON1/CCZ1 are conserved in mammals. **A.** Enlarged Rab5 (magenta) and Rab7 (green) vesicles in *HRS* KO and *CHMP6* KO HeLa cells stably expressing mApple-Rab5 and GFP-Rab7. White arrowheads marking RAB5 and RAB7 positive structures, respectively in the close ups of the individual channels. Yellow arrowheads marking colocalization events in the merge insets. **B.** Quantification of data in (**A**). The size of Rab5 (magenta) positive structures, shown in micron^2^, increases in both of the knockouts compared to control knockout. Each data point represents an average area of one ROI. Obtained *P*-values: Control KO vs. *HRS* KO *P* < 0.0001, Control KO vs. *CHMP6* KO *P* < 0.0001. **C.** Slider model, illustrating the effect of *HRS* or *CHMP6* knockout on the recruitment of Rab5 and Rab7 to endosomal structures (**A** and **B**). The depicted WT level corresponds to the Control KO. **D.** *CCZ1* KO causes the formation of enlarged Rab5 (magenta) positive structures in HeLa cells compared to the control knockout cells. Rab5 positive structures are highlighted via white arrowheads in the insets. **E.** Overexpression of GFP-Rab7 rescues the enlarged size of mApple-Rab5 positive structures in *CCZ1* KO cells. Rab5 and Rab7 positive structures, respectively, are highlighted via white arrowheads in the insets of the individual channels. In the insets of the merged images, colocalization events are marked via yellow arrowheads. **F.** Quantification corresponding to (**D**) and (**E**). Reduction in the area of Rab5 positive structure (magenta) when GFP-Rab7 is overexpressed. Each data point represents an average area of one ROI. *P*-values: Rab5 OE (Control KO) vs. Rab5+Rab7 OE (Control KO) *P* = 0.0003, Rab5 OE (Control KO) vs. Rab5 OE (*CCZ1* KO) *P* < 0.0001, Rab5+Rab7 OE (Control KO) vs. Rab5+Rab7 OE (*CCZ1* KO) *P* > 0.9999, Rab5 OE (*CCZ1* KO) vs. Rab5+Rab7 OE (*CCZ1* KO) *P* < 0.0001. **G.** Slider model, showing the effect of Rab5 or Rab7 overexpression on the recruitment of Rab5 and Rab7 to endosomal structures in *CCZ1* KO background (**D**-**F**). The WT level is shown as grey dotted line for comparison. **H.** Overexpression of HRS-RFP rescues the enlarged size of GFP-Rab5 positive structures in *CCZ1* KO cells. In the close ups of the individual channels Rab5 and HRS, respectively, are marked via white arrowheads. Yellow arrowheads highlight colocalization events in the insets of the merged images. **I.** Quantification corresponding to (**H**). Reduction in the area of GFP-Rab5 positive structure upon overexpression of HRS-RFP. Each data point represents an average area of one ROI. *P*-values: EV+Rab5 OE (Control KO) vs. HRS+Rab5 OE (Control KO) P = 0.7955, EV+Rab5 OE (Control KO) vs. EV+Rab5 OE (*CCZ1* KO) P < 0.0001, HRS+Rab5 OE (Control KO) vs. HRS+Rab5 OE (*CCZ1* KO) P = 0.9912, EV+RAB5 OE (*CCZ1* KO) vs. HRS+Rab5 OE (*CCZ1* KO) P < 0.0001. **J.** Slider model, displaying the effect of Rab5 or HRS overexpression on the recruitment of Rab5 on endosomal structures in *CCZ1* KO background (**H** and **I**). The WT level is shown as grey dotted line for comparison. Data information: Merged images were individually adjusted in all panels. Representative pictures with magnifications (white box) on the right upper corner are shown for each condition (scale bars: 10 μm) in (**A**, **D**, **E** and **H**); *n* = 3 independent experiments. For the quantifications shown in (**B**, **F** and **I**) 1500 structures were measured each (*n* = 1500). Kruskal-Wallis test with Dunn’s multiple comparisons test was performed for each experiment to determine the statistical significance between the examined conditions (**B** and **F**). For (**I**) a Brown-Forsythe ANOVA test and Welch’s ANOVA test, followed by Dunnett’s T3 multiple comparisons test were performed for the same purpose. Significance levels are displayed in the graphs as *** *P* ≤ 0.001, **** *P* ≤ 0.0001 and ns *P* > 0.05. The data show the mean ± s.d. for all examined conditions (**B**, **F** and **I**). Unprocessed images and statistical raw data are available as source data.

## Discussion

How different processes during endosome maturation are coordinated is still poorly understood. Here, we investigated the interplay between Rab conversion and intraluminal vesicle (ILV) biogenesis by the ESCRT machinery. We found that early ESCRT components, in particular HRS, act upstream of Rab conversion and that sorting of cargoes destined for degradation is a prerequisite for the Rab5-to-Rab7 transition to occur. Our data indicate that ubiquitinated cargoes have to be sorted away from the place where Rab conversion can occur into domains that will give rise to ILVs on endosomes. This finding was initially surprising because Rab11-dependent sorting to the plasma membrane can occur from both Rab5 and Rab7 positive endosomes (Podinovskaia *et al*., 2021). However, the Rab5GEF Rabex5 binds to ubiquitin and hence as long as ubiquitinated cargo is still present, Rabex5 might be too stably bound to be displaced by the Rab7GEF Mon1/Ccz1 (Dwivedi *et al*., 2011; Lee *et al*., 2006; Mattera *et al*., 2006; Penengo *et al*., 2006; Poteryaev *et al*., 2010). Thus, we propose that successful corralling of ubquitinated cargo is a pre-requisite for Rab conversion. This model is supported by findings in yeast where deletion of either component of Rab7GEF Mon1/Ccz1 is synthetic lethal with loss of the yeast HRS homolog Vps27 (Aguilar *et al*, 2010; Costanzo *et al*, 2010; Costanzo *et al*, 2016; Surma *et al*, 2013). Likewise, Δ*ccz1* and Δ*vps9*, which encodes for yeast Rabex5 are synthetic lethal (Kucharczyk *et al*, 2009).

It seems as if slowly an order of events is emerging that governs endosome maturation. It appears as if sorting ubiquitinated cargoes into domains that will give rise to ILVs is a pre-requisite for the initiation of Rab conversion. Thus, ubquitinated cargo availability may determine the rate at which endosomes can exchange their Rab proteins. We also know that Rab conversion is critical for endosome acidification (Podinovskaia *et al*., 2021). We still do not know what determines that early endosomes stop accepting new incoming cargo and initiate the maturation process (Huotari & Helenius, 2011; Maxfield & McGraw, 2004). Moreover, even though Rab11 dependent recycling can occur throughout the maturation process, it is likely that a signal exists, which indicates that recycling is completed. Such a signal could for example be the loss of the tubular part of the maturing endosomes or the loss of actin or sorting nexins. Also, it is clear that in a number of cellular systems, transport of the endosomes to the cell center is important for the maturation process (Huotari & Helenius, 2011; McDermott & Kim, 2015; Podinovskaia & Spang, 2018; Solinger & Spang, 2022). Again, how this transport is integrated into the overall maturation process still needs to be established.

We find that the HOPS complex, and in particular VPS-39, can recruit RAB-7 onto endosomes in the absence of SAND-1. In yeast, Vps39 was proposed previously to act as Rab7GEF (Wurmser *et al*., 2000). However, this possibility was largely dismissed by the identification of Mon1/Ccz1 as Rab7GEF, which has a higher GEF activity towards Rab7 than the HOPS complex (Nordmann *et al*., 2010). The presence of a second Rab7GEF, however, is able to explain a number of other observations. First, SAND-1 appears concomitant with RAB-7 on endosomes, but while SAND-1 disappears after about 2 min during Rab conversion, RAB-7 stays behind and its levels even increase afterwards (Poteryaev *et al*., 2010). Thus, Mon1/Ccz1 acts as the switch through coincidence detection of Rabex-5 and PI3P, while HOPS activates Rab7 in a more sustainable manner. Moreover, the existence of a second Rab7GEF can also explain the observed bypass of *sand-1(KO)* caused by *vps-33.1+2(RNAi)* and the partial rescue of the Rab conversion deficiency in knockdowns of the FERARI subunits *SPE-39* and *VPS-45* (Solinger *et al*., 2020; Solinger & Spang, 2014). Finally, Mon1/Ccz1 binds PI3P but does not bind PI3,5P_2_ efficiently (Poteryaev *et al*., 2010). It is tempting to speculate that similar events take place during autophagosome maturation before fusion with lysosomes.

## Materials and Methods

### Worm husbandry and general methods

*C. elegans* were grown and crossed according to standard methods (Brenner, 1974). All experiments and strains were grown at 20 °C. At this temperature *sand-1(KO)* strains are still viable but show already severe endosomal trafficking phenotypes (Poteryaev *et al*., 2010; Poteryaev *et al*., 2007; Solinger & Spang, 2014). The following *C. elegans* transgenes were used in this study: *sand-1(ok1963)IV*, *unc-119(ed3)III*, *pwIs72[pvha6::GFP::rab-5 + unc-119(+)]*, *pwIs846[Pvha-6-RFP-rab-5; Cb.unc-119(+)]*, *pwIs429[pvha-6::mCherry::rab-7]*, *pwIs90[Pvha-6::hTfR-GFP; Cbr-unc-119(+)]*, *pwIs50[Plmp-1::lmp-1::GFP + Cb-unc-119(+)]*, *pwIs518[vha-6::GFP-HGRS-1]* and *jyEx128 [vha-6p:::GFP::UBQ, cb-unc-119(+)]*. Strains used in this study are listed in Appendix Table S9.

### Gene knockdowns in *C. elegans*

Knockdowns were performed via feeding (Solinger *et al*, 2014). For each experiment 5 L4/young adult worms (F0) were transferred on isopropyl β-D-1-thiogalactopyranoside (IPTG) containing standard nematode growth medium (NGM) agar plate seeded with dsRNA producing bacteria. In case of pre-feeding experiments, several worms (F0) were transferred first on IPTG plates seeded with control dsRNAi producing bacteria. L3 worms of the F1 generation were transferred on plates for RNAi. Several plates per condition were set up in parallel as technical replicates, and F1 worms were imaged when they reached young adult stage in all experiments. RNAi constructs expressing plasmids transformed in *E. coli* HT115 used for feeding were obtained from Ahringer library (Kamath *et al*, 2003) or created via cloning (see plasmid generation for *C. elegans*). All used constructs were confirmed via sequencing (Microsynth AG, Balgach, Switzerland).

### Microscopy of *C. elegans*

Worms were imaged with a confocal laser scanning microscope Olympus FV3000 (Olympus Schweiz AG, Wallisellen, Switzerland) unless stated otherwise. For imaging following lasers were used with intensities between 0.8-10%: 405 nm, 488 nm and 561 nm. To obtain high resolution pictures the Galvano scan device was used. The sampling speed was set to 8.0 μs/pixel and the scan direction was, one way scanning, during imaging. Pictures were taken with a UPLSAPO60XS2 objective lens and silicone immersion oil (Olympus Schweiz AG). For image acquisition and processing at the microscope FV31S-SW software (Olympus Corp, Tokyo, Japan) was used. For imaging, worms were transferred from the plate in a drop of Levamisole (0.5 ml Levamisole (100 mM) + 1 ml M9) on a 2% agarose pad on a slide. The coverslip was sealed with Vaseline.

The images of the *rabx-5* knockdown in *sand-1(KO)* were obtained on a Zeiss LSM 880 with Airyscan (Carl Zeiss AG, Feldbach, Switzerland), using 488 nm and 555 nm lasers. The color channels were imaged in separate phases to reduce bleed through. All images were taken and processed at the microscope using the Zen Blue software (Carl Zeiss AG, Oberkochen, Germany). Worms were paralyzed with 20 mM NaN_3_.

### Transmission electron microscopy (TEM)

For TEM, worms were frozen as follows*. C. elegans* animals were picked with a worm pick from an agar plate and transferred to a droplet of M9 medium on a 100 µm cavity of a 3 mm aluminium specimen carrier (Engineering office M. Wohlwend GmbH, Sennwald, Switzerland). 5 - 10 worms were added to the droplet and the excess M9 medium was sucked off with dental filter tips. A flat aluminium specimen carrier was dipped in 1-hexadecene and added on top. Immediately, the specimen carrier sandwich was transferred to the middle plate of an EM HPM100 high-pressure freezer (Leica Microsystems GmbH, Vienna, Austria) and frozen immediately.

Freeze-substitution was carried out in an AFS2 freeze-substitution unit (Leica Microsystems GmbH) with integrated, custom built agitation system in water-free acetone containing 1% OsO_4_ for 8 hr at -90°C, 7 hr at -60°C, 5 hr at -30°C, 1 hr at 0°C, with transition gradients of 30°C/hr, followed by 30 min incubation at RT. Samples were rinsed twice with acetone water-free, block-stained with 1% uranyl acetate in acetone (stock solution: 20% in methanol) for 1 hr at 4°C, rinsed twice with water-free acetone and embedded in Epon-Araldite (Merck KGaA, Darmstadt, Germany) (exact composition visible in Appendix method section): 66% in acetone overnight, 100% for 1 hr at RT and polymerized at 60°C for 28 hrs. Ultrathin sections (50 nm) were transferred on one hole grids (with a Formvar film and coated with 15 nm of carbon) and post-stained with Reynold’s lead citrate (exact protocol visible in Appendix method section) and imaged in a Talos 120 transmission electron microscope at 120 kV acceleration voltage equipped with a bottom mounted Ceta camera (Thermo Fisher Scientific FEI Europe B.V., Eindhoven, Netherlands) using the Maps software (Thermo Fisher Scientific Inc., Waltham, USA). Image analysis was performed via Maps software (Thermo Fisher Scientific Inc.), Fiji and Prism (GraphPad Software LLC., Boston, USA)

### *C. elegans* MVB and ILV quantifications

To quantify the MVB and ILV phenotypes of *C. elegans* intestinal cells, unstitched high resolution (original pixel size = 8.2169 Å) TEM tile sets showing ultrathin sections of whole worm guts were examined via Maps software (Thermo Fisher Scientific Inc.) to detect all MVBs (per worm one section was analyzed). In the next step tiles showing MVBs were taken and the in this way detected MVBs were analyzed regarding their size, ILV number and ILV size via Fiji. So, for each MVB in the examined guts the corresponding number of ILVs, the size of the ILVs and the size of the MVB itself were measured manually (see Appendix Fig. S2). Results of these measurements were than transferred into Excel (Microsoft Corp. Redmond, USA) for further calculations or directly in Prism (GraphPad Software LLC.) and finally statistically analyzed and visualized via Prism (GraphPad Software LLC.).

### *C. elegans* population growth analysis

The growth of the worm populations during the RNAi experiments was evaluated via binocular microscopy using a Stemi 2000 stereomicroscope equipped with a KL 200 light source (Carl Zeiss AG). After several days of incubation, when the F1 worms of the control group reached young adult stage, the populations were examined and the knockdowns were compared with the control group. Each experiment was performed independently three times.

### Area localization quantification of RAB-7 and LMP-1 in *C. elegans*

The area localization quantification of the marked proteins in the intestine was performed manually via Fiji. To get comparable results, only worm slices which showing the middle plane of the gut (z-dimension) were used for the analysis. Per worm one slice from the z-stack was isolated and analyzed. This picture was then rotated in a way that the worm gut was shifted perpendicular. To measure the area localization a ROI (200 × 650 pixel) was placed in the worm on a representative spot covering gut and lumen horizontal. Afterwards the scale of the picture was deleted to get a result in pixels and a line measurement using “plot profile” was performed. This measurement generates a line plot consisting of 650 values describing the average brightness of a 200-pixel long line each (see Fig. EV4A). The values were then transferred into Excel (Microsoft Corp.). In case of a multichannel image the same ROI was placed on the equivalent spot in the second channel and the measurement was repeated in the same way. The same ROI was used for each worm throughout the whole quantification to be consistent. In Excel (Microsoft Corp.) the brightness values of each worm were normalized to 100. The final mean values were again normalized to 100 to obtain comparable values and graphs.

### Quantification of number and size of RAB-5 or hTfR positive endosomes in *C. elegans*

The size of RAB-5 or hTfR positive structures in the worm gut was semi automatized quantified using Fiji. Each worm was treated in the following way. For analysis one slice of the z-stack showing the middle plane of the worm gut (z-dimension) was identified and a threshold was set to highlight distinct structures and reduce the background. Afterwards the picture is converted into binary and treated with the “open” and “watershed” functions to further reduce background and sharpen the structures. The remaining structures are then measured via the “analyze particles” function to obtain the area value of each particle. The obtained results were then transferred into Excel (Microsoft Corp.) and Prism (GraphPad Software LLC.) for further analyzes and following graphical depiction. To analyze the number of RAB-5 or hTfR positive structures in the worm gut the number of RAB-5 or hTfR particles of each worm used to determine the RAB-5 or hTfR particle size was counted and correlated with the gut area visible on the analyzed pictures. Statistical analyzes and plotting were performed in Prism (GraphPad Software LLC.).

### Colocalization examination of RAB-5, RAB-7, HGRS-1 and LMP-1

Colocalization of two markers in *C. elegans* intestinal cells was quantified via Fiji. For each analyzed worm one slice of the captured z-stack, showing the middle plane, was taken out and the two relevant channels were aligned manually or computer assisted (Fiji plugins TurboReg and MultiStackReg (see Appendix Table S14)). In these pictures a ROI (300 × 400) spanning one half of the gut was drawn and the colocalization of the two markers in these ROIs were measured (see Fig. 3B) via the Fiji plugin JaCop (see Appendix Table S14). The thresholds to calculate the Mander’s coefficients (M1 and M2) were adjusted in each picture and for each channel individually to measure representative signals for each marker. Statistical analyzes and visualization were performed in Prism (GraphPad Software LLC.).

### Quantification of UBQ localization and structure size in *C. elegans*

The nuclear UBQ accumulation in comparison to the cytoplasmic UBQ abundance and the size of UBQ-positive structures in *C. elegans* intestinal cells were measured manually using Fiji.

The localization of the tagged UBQ was measured via ROI (50 × 50) in the following way. In the selected plane each nucleus and a cytoplasmic region next to them were analyzed for their raw fluorescence intensity using the same ROI. The obtained values were transferred into Excel (Microsoft Corp.) and used to calculate the change of fluorescence intensity between nucleus and cytoplasm for each nucleus. To visualize the results and perform statistical analyzes the calculated changes were transferred into Prism (GraphPad Software LLC.).

The size of the UBQ positive structures was measured in the selected plane with the “analyze particles” function, like described in “quantification of number and size of RAB-5 positive endosomes in *C. elegans*”. However, here an extra step was performed before the thresholding. To avoid counting bright nuclei as cytoplasmic UBQ positive structures the nuclei were circled manually and cut out. The in this way measured particle sizes were then transferred into Excel (Microsoft Corp.) and finally into Prism (GraphPad Software LLC.) for statistical analyzes and visualization.

### Cell culture, transfection procedure and CRISPR-Cas9 KO in mammalian cells

HeLa CCL2 cells were grown at 37°C and 5% CO_2_ in high-glucose Dulbecco’s modified Eagle’s medium (DMEM) (Sigma-Aldrich), supplemented with 10% fetal calf serum (FCS) (Biowest S.A.S., Nuaillé, France), 2 mM L-Glutamine (Gibco (Thermo Fisher Scientific Inc.)), 1 mM Sodium Pyruvate (Sigma-Aldrich), and 1 x Penicillin and Streptomycin (Sigma-Aldrich). The HeLa cell line was a kind gift of Martin Spiess and its identity was authenticated by short tandem repeat (STR) analysis from an external service provider (Microsynth AG). All cell lines created and used for the study were confirmed to be mycoplasma-negative via PCR and are listed in Appendix Table S10.

For transient cell transfections, cells were plated into 6-well plates to reach 70% confluency the following day and transfected with 1µg plasmid DNA complexed with Helix-IN transfection reagent (OZ Biosciences S.A.S., Marseille, France).

For CRISPR/Cas9-mediated knockout, guide RNAs were selected using the CRISPR design tool CHOPCHOP (see Appendix Table S14). Two guide RNAs were designed from two different exons for each target gene (*HRS*: exon 1 and exon 22; *CHMP6*: exon 3 and exon 5). Annealed oligonucleotides were cloned into two different plasmids containing mTagBFP2 marker and Puromycin resistance, respectively (Px458 mTagBFP2 and Px459 Puro). For primer sequences see Appendix Table S11. Constructs were verified by sequencing (Microsynth AG). In brief, HeLa cells stably expressing mApple-Rab5 and GFP-Rab7 were seeded at 2□×□10^6^ cells per 10□cm dish. The following day, cells were transfected with 2.5□μg of the plasmids (control vectors without insert or vectors containing a guide RNA against target gene). Transfecting media was exchanged with fresh media after 6 h. Cells were treated with puromycin for 24 h after transfection followed by fluorescence-activated cell sorting (FACS) on next day. For FACS, 48□h after transfection, cells were trypsinized and resuspended in cell-sorting medium (2% FCS and 2.5 mM ethylenediaminetetraacetic acid (EDTA) in phosphate-buffered saline (PBS)) and sorted on BD FACS AriaIII Cell Sorter (Becton, Dickinson and Company Corp., Franklin Lakes, USA). GFP, mApple and BFP positive cells were collected and seeded in a new well.

### Live cell imaging of mammalian cells

For live imaging, cells were plated in 8 well chambered coverglass and media was replaced with warm imaging buffer (5 mM dextrose (D(+)-glucose, H2O, 1 mM CaCl2, 2.7 mM KCl, 0.5 mM MgCl2 in PBS) just before imaging. Images were taken at 37°C on an inverted Axio Observer Zeiss microscope (Carl Zeiss AG) equipped with a TempModule S, a Heating Unit XL S, an incubator XLmulti S1 (PeCon GmbH, Erbach, Germany), a HXP 120 V (Leistungselektronik JENA GmbH, Jena, Germany) and a SMC 2009 (Carl Zeiss AG). For imaging a Plan Apochromat N 63×/1.40 oil DIC M27 objective with a Photometrics Prime 95B camera (Teledyne Photometrics Inc., Tucson, USA) was used. Z-stack images were captured using the Zen Blue 2.6 imaging software (Carl Zeiss AG) and deconvolved using Huygens Remote Manager (HRM) (Huygens Software (Scientific Volume Imaging B.V., Hilversum, Netherlands)). Acquired images were processed by using the OMERO client server web tool and Fiji.

### Rab5 structure size quantification in mammalian cells

Area analysis for Rab5 positive structures was done using “analyze particle” function of Fiji. To elaborate more, initially, Rab5 and Rab7 channels were separated using split channels function of Fiji. Image was converted to the maximum projection using z stacks acquired during imaging. Further, a ROI of 100 μm^2^ was drawn manually and duplicated. Objects were manually selected using threshold based on the intensity. Areas of individual Rab5 positive structures in the ROI were acquired using the analyze particle function. The average area of Rab5 positive particles in one ROI was plotted using Prism (GraphPad Software LLC.).

### Plasmid construction and source for experiments and knockouts in mammalian cells

All constructs used in this study are listed in the Appendix Table S12.

### Plasmid construction for *C. elegans* experiments

To generate the RNAi constructs for knockdowns of *tsg-101*, *vps-2*, *vps-60* and *rabx-5*, primers with homology overhangs for pDT7 (L4440) (Timmons and Fire, 1998, Timmons et al., 2001) were designed via NEBuilder Assembly Tool (New England Biolabs Inc., Ipswich, USA) (see Appendix Table S13). Using these primers, the corresponding sequences from cDNA or genomic DNA of *C. elegans* were amplified via PCR. The in this way generated constructs were afterwards incorporated in pDT7 at the EcoRV cutting site via standard Gibson assembly. The Gibson assemblies were then transformed in *E. coli* and the generated plasmids were verified by sequencing (Microsynth AG).

### Statistical analysis

Statistical analyses were performed using Prism (GraphPad Software LLC.). Used statistical tests and the determined *P*-values (in case of *P* = 0.0001 with four decimal places, two extra decimal places are given to clarify the value) are indicated in the legends belonging to the depicted graphs. The data sets were tested for normality/lognormality with Anderson-Darling test, Shapiro-Wilk test, Kolmogorov-Smirnov test and D’Agostino & Pearson test, for differences in the s.d. with Brown-Forsythe test and Bartlett’s test and for differences in the variance with F test. For each test a significance level (alpha) of 0.05 was used. All programs used in this study are listed in the appendix (Appendix Table S14).

## Acknowledgements

We thank Barth D. Grant, Emily R. Troemel, Attila Stetak, and Martin Spiess for worm strains and HeLa cells, respectively. Some strains were provided by the CGC, which is funded by the NIH Office for Research Infrastructure Programs (P40 OG010440). We acknowledge Dora Stetak for technical support and Julia K. Nussbaum and Arian Dill for their help in plasmid construction. The imaging core facility of the Biozentrum (IMCF) is acknowledged for superb support. We thank Carmen Kaiser from the Center for Microscopy and Image Analysis, University of Zurich for ultra-thin sectioning of embedded *C. elegans* for TEM. Cells were sorted in the Biozentrum FACS core facility. This work was supported by the University of Basel and the Swiss National Science Foundation (310030_197779, 310030_185127).

## Disclosure and competing interests statement

The authors declare no competing interests.

## Expanded view figure legends

**Figure 1: Knockdowns of core early and late ESCRT factors as well as knockdowns of additional ESCRT-III factors affect endosome maturation in WT and *sand-1(KO)*, related to Figure 1**.

**A.** Additional ESCRT knockdowns performed in the ESCRT RNAi screen shown in Fig. 1A. White arrowheads pointing to GFP::RAB-5 and mCherry::RAB-7 positive structures, respectively in the individual channels. In the merges colocalization events are marked via yellow arrowheads (signals: RAB-5 > green and RAB-7 > magenta).
**B.** Further ESCRT knockdowns performed in the ESCRT RNAi screen shown in Fig. 1B. Consistent with data shown in Fig. 1B, knockdowns of further core early ESCRTs or a core late ESCRT factor cause no further enlargement of the through *sand-1(KO)* enlarged GFP::RAB-5 positive structures and have only minor effects on the colocalization of GFP::RAB-5 and mCherry::RAB-7. Similar effects on the colocalization of GFP::RAB-5 and mCherry::RAB-7 and the GFP::RAB-5 structure size are also observable in knockdowns of additional ESCRT-III factors. White arrowheads pointing to GFP::RAB-5 and mCherry::RAB-7 positive structures, respectively in the individual channels. In the merges colocalization events are marked via yellow arrowheads (signals: RAB-5 > green and RAB-7 > magenta).

Data information: Merges were individually adjusted in all panels. Representative pictures with magnifications (white box) on the right are shown for each experiment (scale bars: 10 μm (main pictures) and 1 μm (magnifications)). Unprocessed images are available as source data.

**Figure 2: ESCRT knockdowns affect the intracellular morphology in WT and *sand-1(KO)*, related to Figure 2**.

**A.** TEM overview pictures belonging to experiments shown in (Fig. 2 **A**, **C** and **E**). The *sand-1(KO)* causes the formation of large granular structures accumulating in the basal region but does not increase the MVB size. Knockdown of *tsg-101* increases the size of endosomal structures, whereas *vps-2* knockdown causes no major changes in this regard but leads to the formation of additional small and big granular structures. The observed ESCRT RNAi effect are strain background independent.

Data information: Representative overview TEM pictures are shown for each examined condition. Overview electron micrographs are individually adjusted and were generated with unstitched data sets (scale bars 10 μm). Unprocessed images are available as source data.

**Figure 3: *usp-50* and *did-2* knockdown cause UBQ aggregation in the cytoplasm and USP-50 presence is required for HGRS-1 stability in *C. elegans*, related to Figure 4**.

**A.** The knockdown of *did-2* has only minor effects on nuclear GFP::UBQ levels and cause the formation of GFP::UBQ accumulations in addition to enlarged structures. *usp-50(RNAi)* however, generates GFP::UBQ accumulations, enlarged structures, tubular networks and strongly reduces nuclear GFP::UBQ levels. White arrowheads pointing to GFP::UBQ positive structures and corresponding areas in the DIC, respectively.
**B.** Early and late ESCRT knockdowns showing with equal brightness settings, corresponding to Fig. 4. *hgrs-1(RNAi)* and *vps-20(RNAi)* show a strong reduced GFP::UBQ level in comparison to Mock as well as to the other ESCRT knockdowns.
**C.** Quantification of the GFP::UBQ positive structure sizes belonging to (**A**) and Fig. 4A. The individual sizes of the measured GFP::UBQ positive structures are displayed for each examined condition in micron^2^. For analysis shown in the graph 10 worms per condition (*n* = 10) and more than 9500 structures in total (*n* > 9500) were examined (*n* = 3 independent experiments). The mean particle size is shown for each condition as red line in the graph and the data are shown in log 10 scale (Y-axis). Obtained *P*-values: Mock vs. *hgrs-1(RNAi) P* < 0.0001, Mock vs. *tsg-101(RNAi) P* = 0.3887, Mock vs. *vps-20(RNAi) P* > 0.9999, Mock vs. *vps-24(RNAi) P* < 0.0001, Mock vs. *vps-2(RNAi) P* = 0.1113, Mock vs. *did-2(RNAi) P* < 0.0001, Mock vs. *vps-4(RNAi) P* < 0.0001, Mock vs. *usp-50(RNAi) P* < 0.0001.
**D.** The knockdown of *usp-50* causes a strong reduction of the GFP::HGRS-1 signal and generates tubular, network like RFP::RAB-5 positive structures and RFP::RAB-5 positive aggregates. In addition, some of the remaining GFP::HGRS-1 positive structures are also positive for RFP::RAB-5. White arrowheads marking GFP::HGRS-1 and RFP::RAB-5 positive structures, respectively in the individual channels. Yellow arrowheads indicating colocalization events in the merges (Signals: HGRS-1 > green and RAB-5 > magenta).

Data information: Representative pictures with corresponding enlargements (white box) next to it are shown for each experiment (scale bars: 10 μm (main pictures) and 1 μm (enlargements)) in (**A**, **B** and **D**). Belonging DIC pictures elucidate the extent of the gut, marker independent are depicted for all experiments showing in (**A**). Kruskal-Wallis test with Dunn’s multiple comparisons test was performed to determine the statistical significance between the examined conditions (**C**). Significance levels are displayed in the graph as **** *P* ≤ 0.0001. Comparisons with *P* > 0.05 are not shown for simplification. Unprocessed images and statistical raw data are available as source data.

**Figure 4: LMP-1 and RAB-7 area localizations in *sand-1(KO)* under reduced UBQ levels and knockdowns of *vps-39*, related to Figures 5 and 6**.

**A.** Schematic representation of the approach used to generate the area plots shown in Fig. 5, 6 and EV4. A simplified worm gut with intestinal cells expressing a blue example protein is shown on the left side. The localization of this protein gets measured via a ROI (200 x 650 pixels) spanning the whole worm gut. This creates an area plot, highlighting the localization of the protein in the intestinal cells, shown on the right side.
**B,C**. Area plots belonging to Fig. 5I and K. Overexpression of mCherry::RAB-7 causes an apical localization of LMP-1::GFP which can be further augmented through RNAi mediated UBQ level reduction.
**D,E**. Area plots belonging to Fig. 6C and D. The knockdown of HOPS subunit *vps-39* abolishes the partial rescue of *sand-1(KO)* mediated by the mCherry::RAB-7 overexpression and causes a more basal distribution of LMP-1::GFP. This localization shift is less prominent if the UBQ levels are reduced together with the *vps-39* knockdown.
**F**. Examined mCherry::RAB-7 area localization belonging to Fig. 6A. The knockdown of *vps-39* impairs the recruitment of mCherry::RAB-7 to apical structures caused by the reduction of UBQ levels. The control group is the same like shown in Fig. 5B and the data were collected and analyzed like described for this experiment.

Data information: The individual mCherry::RAB-7 area plots for the merge area plots are shown in Fig. 5K and 6D. Statistical raw data are available as source data.

**Figure 5: UBQ reduction reduces the number of RAB-5 structures and *usp-50(RNAi)* causes colocalization of RAB-5 with RAB-7 and UBQ.**

**A.** The reduction of the UBQ level via RNAi causes a depletion of GFP::RAB-5 positive structures. Knockdown of *usp-50* causes a mislocalization of GFP::RAB and mCherry::RAB-7 and increases their colocalization. White arrowheads marking GFP::RAB-5 and mCherry::RAB-7 positive structures, respectively in the individual channels. Yellow arrowheads marking colocalization events in the merges (Signals: RAB-5 > green and RAB-7 > magenta).
**B.** Schematic representation of the endosomal RAB-5/RAB-7 balance in a *C. elegans* intestinal cell in WT background. Effects of UBQ reduction or *usp-50(RNAi)* on this balance are shown via black arrows.
**C.** The RNAi mediated UBQ level abatement causes the formation of fewer and less prominent RFP::RAB-5 aggregates and leads to fewer GFP::UBQ structures. *usp-50(RNAi)* causes the formation of large GFP::UBQ structures and aggregates which are often also RFP::RAB-5 positive and reduces the GFP::UBQ signal in the nucleus. White arrowheads marking GFP::UBQ and RFP::RAB-5 positive structures, respectively in the individual channels. Yellow arrowheads marking colocalization events in the merges (Signals: UBQ > green and RAB-5 > magenta).
**D.** Schematic representation of the UBQ distribution in an intestinal *C. elegans* cell expressing marked UBQ and RAB-5 in WT background. The effect of the *usp-50* knockdown on this equilibrium is shown via a black arrow.

Data information: Merges were individually adjusted in all panels. Representative pictures with close ups (white box) on the right are shown for each experiment (scale bars: 10 μm (main pictures) and 1 μm (close ups) in (**A** and **C**); *n* = 3 independent experiments. Unprocessed images are available as source data.

## Notes

### Competing Interest Statement

The authors have declared no competing interest.

